# The EGF/EGFR-YAP1/TEAD2 axis-mediated BRAF inhibitor vemurafenib resistance upregulates STIM1 in melanoma

**DOI:** 10.1101/2024.01.29.577717

**Authors:** Weiyu Bai, Chenghao Yan, Yichen Yang, Lei Sang, Qinggang Hao, Xinyi Yao, Jia Yu, Yifan Wang, Xiaowen Li, Mingyao Meng, Jilong Yang, Junling Shen, Yan Sun, Jianwei Sun

**Affiliations:** Center for Life Sciences, Yunnan Key Laboratory of Cell Metabolism and Diseases, State Key Laboratory for Conservation and Utilization of Bio-Resources in Yunnan, School of Life Sciences, Yunnan University, Kunming, 650091, China; Tianjin Medical University Cancer Institute and Hospital. National Clinical Research Center for Cancer. Key Laboratory of Cancer Prevention and Therapy, Tianjin’s Clinical Research Center for Cancer. Tianjin, China; Key Laboratory of Tumor Immunological Prevention and Treatment of Yunnan Province, Kunming, China

**Keywords:** STIM1, EGF/EGFR, YAP1, Melanoma, BRAF inhibition

## Abstract

STIM1 is the endoplasmic reticulum (ER) Ca^2+^ sensor for store-operated entry (SOCE) and closely correlated to carcinogenesis and tumor progression. Previously we found that STIM1 is upregulated in melanoma cells resistant to the BRAF inhibitor vemurafenib, but the regulation mechanism is unknown. Here, we show that vemurafenib resistance upregulates STIM1 through an EGF/EGFR-YAP1/TEAD2 axis. Vemurafenib resistance can lead to the increase of EGF and EGFR levels to activate the EGFR signaling pathway. Reactivated EGFR signal promotes YAP1 nuclear localization to increase the expression of STIM1. Our finding not only demonstrates the mechanism by which vemurafenib resistance promotes STIM1 expression, but also provides combined targeting EGF/EGFR-YAP1/TEAD2-STIM1 to improve the therapeutic efficiency of BRAF inhibitor in melanoma patients.

## Introduction

Melanoma, a highly malignant tumor derived from the malignant transformation of melanocytes. The incidence rate and mortality of malignant melanoma has been increasing in recent decades. Mutational activation *BRAF* is the most prevalent genetic alteration in human melanoma, with more than 50% of the patients harboring the *BRAF(V600E)* mutation[1–3]. Several mutant *BRAF* specific inhibitors (BRAFi) such as vemurafenib have been developed to specifically inhibit *BRAF(V600E)* mutations and remarkably improved the survival of melanoma patients with oncogenic BRAF mutations. However, despite the rapid progress in *BRAF*-mutant melanoma treatment, drug resistance is still the main obstacles for vemurafenib therapy and treatment.

Vemurafenib resistance occurred in about 50% patients and half patients had disease progression within 6 months from the start of vemurafenib monotherapy, which greatly hinders the clinical application of vemurafenib. Therefore, exploring *BRAF(V600E)*-associated prognostic factors for providing potential joint targets is important for combined targeted therapy.

It has been reported that the re-activation of the MAPK pathway is of particular importance to the resistance to BRAFi therapy[4, 5]. However, an increasing number of studies have shown that the EGF/EGFR pathway is activated in response to vemurafenib treatment in vemurafenib resistant melanoma cells [6, 7].

The epidermal growth factor receptor (EGFR) is a tyrosine kinase receptor and links extracellular signals to control cell survival, growth, proliferation and differentiation. EGFR has been an effective therapeutic target for cancer malignancy due to its frequent mutation and overexpression[8]. Recent studies indicated that intrinsic vemurafenib resistance is due to the feedback activation of EGFR signaling pathway[6, 9, 10]. EGFR-mediated activation of RAS and CRAF then reactivates the MAPK / ERK pathway[10]. Although there are many studies on tumor cell response to BRAF inhibitor resistance, it is still a challenge to clarify how melanoma cells acquired resistance that mediated tumor metastasis.

STIM1 is the key endoplasmic reticulum (ER) Ca^2+^ sensor for store-operated calcium entry (SOCE). We previously reported that SOCE promotes melanoma invasion and metastasis (citation). The STIM1 up-regulation is usually closely related to tumorigenesis and progression [11, 12]. However, the regulation of STIM1 in melanoma and the roles of STIM1 in BRAFI vemurafenib resistance is largely unknown. Ziyi Yang etc reported that STIM1 overexpression was related to the acquired resistance to imatinib[13]. Our recent results show that STIM1 is up-regulated in vemurafenib resistant melanoma cells[14], suggesting that the high expression of STIM1 may respond to BRAF inhibitor resistance in melanoma. However, the mechanism by which vemurafenib induced STIM1 up-regulation is still not completely understood.

Here in this study, we demonstrated that the EGF/EGFR-YAP1/TEAD2 axis mediated STIM1 upregulation in vemurafenib resistant melanoma cells. Vemurafenib resistance can lead to the increase of EGF and EGFR levels and activate the EGFR signaling pathway, which promotes the nuclear translocation of YAP1 and activates the Hippo pathway, YAP1/TEAD2 bind to the promoter of STIM1 and promote the transcription of STIM1. Our study indicated that the EGF/EGFR-YAP1-STIM1 axis is a potential combined therapeutic target with BRAFi therapy for melanoma. The present study not only demonstrated the mechanism by which vemurafenib resistance promotes STIM1 expression, but also provided targeting EGF/EGFR-YAP1/TEAD2-STIM1 to improve the therapeutic efficiency of BRAF inhibitor in melanoma patients.

## Result

### 1. STIM1 is upregulated in BRAF inhibitor vemurafenib resistant cells

We have previously reported that the STIM1-PYK2-invadopodia axis mediated vemurafenib-induced metastasis in melanoma[14, 15], Vemurafenib treatment significantly increases *STIM1* mRNA and protein levels in WM793 cells (Fig.1A, B). The level of STIM1 decreased after removing the vemurafenib, which indicates that BRAF inhibition induced STIM1 upregulation. To confirm the hypothesis, we test edthe STIM1 level in WM115 control and vemurafenib resistance cells. Our result indicated BRAF inhibition may increase STIM1 levels (Fig.1C, D). To investigate if BRAF inhibition with shRNA increased STIM1 levels, we knocked down BRAF in WM793 cells, and *BRAF* shRNA could also increase *STIM1* mRNA and protein levels (Fig.1E, F). Our results suggested that inhibition of BRAF may regulate the transcription of STIM1 through the activation of a certain pathway.

**Figure 1.**
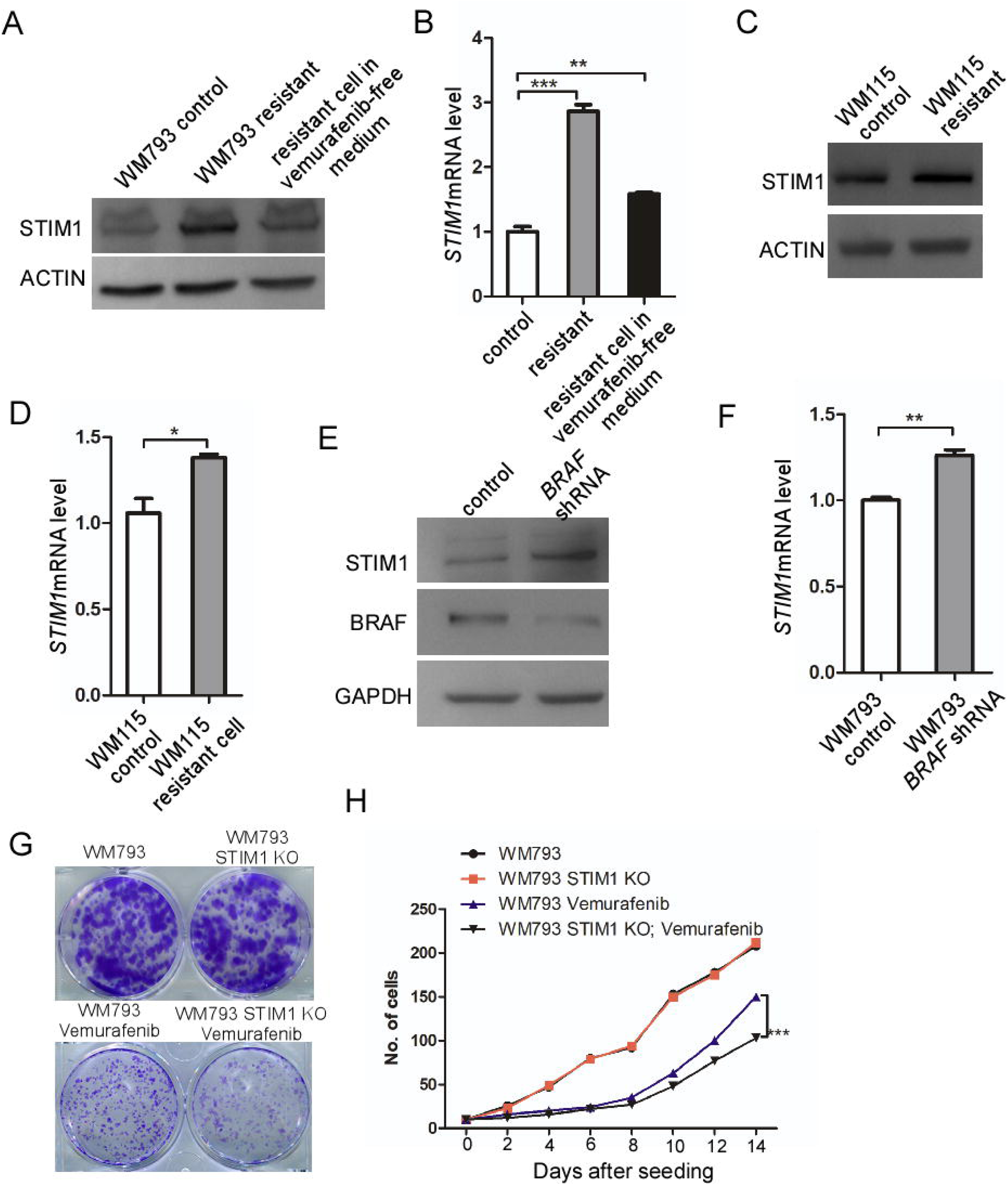
**BRAF inhibition induced STIM1 up-regulation** A. WB analysis of STIM1 level in control and vemurafenib resistant WM793 cells B. Q-PCR analysis of *STIM*1 level in control and vemurafenib resistant WM793 cells C. WB analysis of STIM1 level in control and vemurafenib resistant WM115 cells D. Q-PCR analysis of *STIM*1 level in control and vemurafenib resistant WM115 cells E. WB analysis of STIM1 level in control and *BRAF* shRNA WM793 cells F. Q-PCR analysis of *STIM*1 level in control and *BRAF* shRNA WM793 cells G. The effect of STIM1 on clone formation in presence of vemurafenib H. The effect of STIM1 on cell proliferation in presence of vemurafenib

To investigate if STIM1 is involved in vemurafenib resistance, we knocked out STIM1 in WM793 cells and treated with vemurafenib. And we found that *STIM1* knockout did not affect cell proliferation, but significantly inhibited/reduced cell proliferation in the presence of vemurafenib (Fig.1G, H), which suggested that STIM1 is involved in vemurafenib resistance.

### 2. BRAF inhibition induced EGF/EGFR up-regulation

In order to investigate the mechanism how BRAF inhibition resistance promoted STIM1 expression, we first subject the control, vemurafenib-treated and vemurafenib resistance WM793 cells for RNA sequencing. The result showed that many gene levels were significantly changed in vemurafenib treated and resistant cells. The mRNA levels of 70 genes increased significantly, and 80 genes decreased significantly in vemurafenib treated and resistant cells (Fig. 2A). The upregulated genes are closely related to the Wnt signaling pathway, tumor metastasis and cytoskeleton. Including Hippo and MAPK pathways (Fig. 2C, D), suggesting that the drug resistance caused by BRAF inhibitor treatment can promote tumor metastasis.

**Fig 2:**
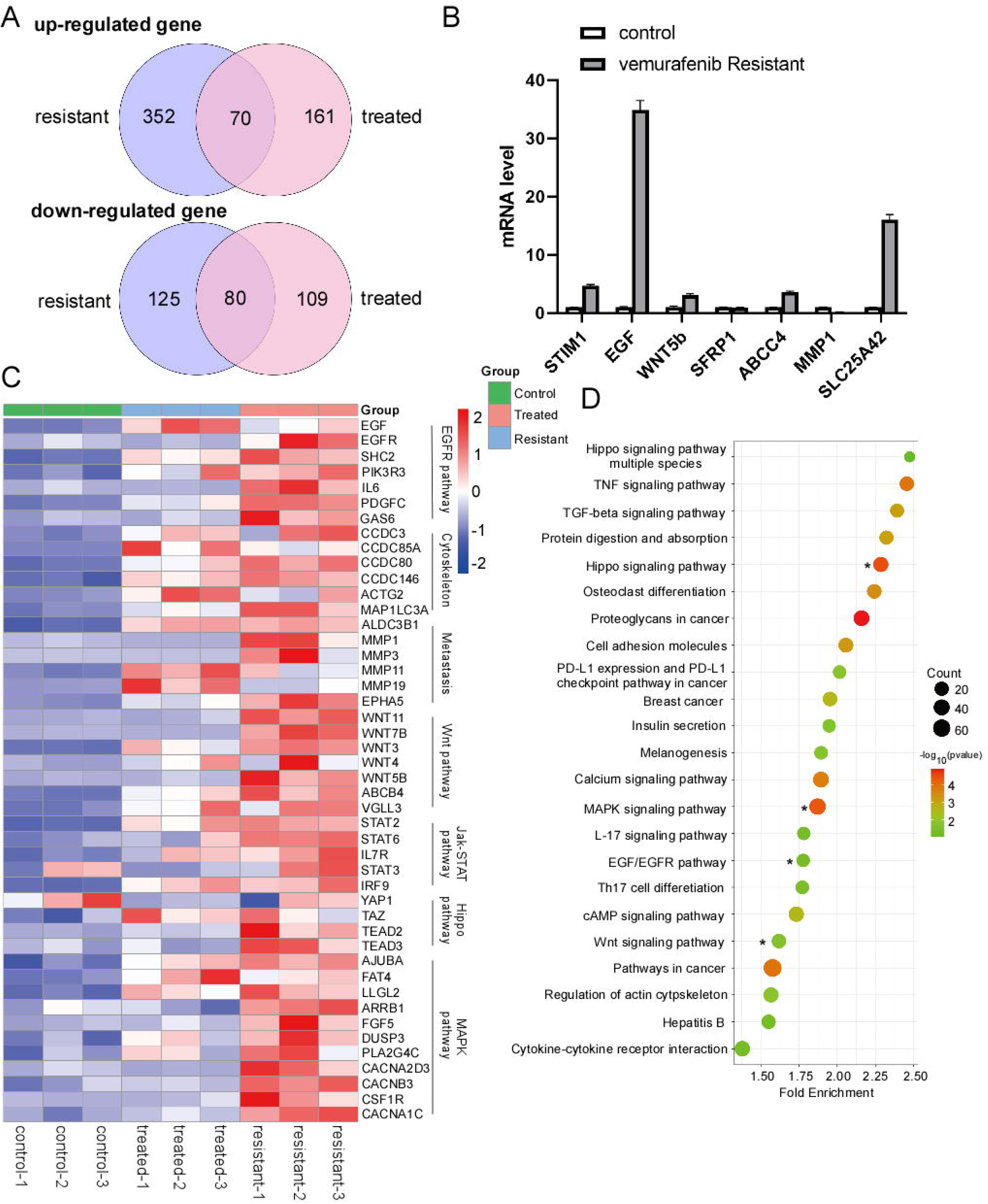
Vemurafenib resistance activates the EGF/EGFR signaling pathway A. Statistics of RNA sequencing results in control and vemurafenib resistant WM793 cells B. Q-PCR analysis of selected genes level in control and vemurafenib resistant WM793 cells C. Heatmap analysis of genes expression in different signaling pathways D. KEGG analysis of differentially expressed genes between control and vemurafenib resistant cells

In order to verify our results, we first selected several genes through qPCR analysis. Our analysis showed that the mRNA level of many secretory proteins was significantly increased in vemurafenib resistant cells, the most obvious of which was EGF (Fig. 2B). It has been reported that EGFR is induced by BRAF inhibitor resistance[6]. We detected EGFR in control and vemurafenib resistant WM793 cells, although our sequencing results did not screen for EGFR, our results showed that the protein and mRNA levels of *EGFR* were significantly increased in vemurafenib resistant WM793 cells, which is the same as previous reports [6]. The results suggested that the EGF/EGFR signaling pathway is activated after vemurafenib resistance.

We then performed GO enrichment and KEGG analysis. We found that the acquired resistance to vemurafenib correlated with the Hippo pathway, which is consistent with previous reports [16].

### 3. Vemurafenib resistance induces STIM1 up-regulation through EGF/EGFR signaling

To investigate the mechanism that Vemurafenib resistance induces STIM1 up-regulation, we then examined if vemurafenib resistant induces STIM1 expression through EGF/EGFR. To examine this hypothesis, we first detected the level of EGF in the supernatant of control and vemurafenib resistant cells. The result indicated that resistant cells secret more EGF, even if vemurafenib is removed from the culture medium (Fig. 3A). To further investigate whether vemurafenib resistance promotes the level of EGF, we constructed vemurafenib resistant WM115 cell line. Compared to control cells, vemurafenib resistant WM115 cells can also produce more EGF (Fig. 3C). Our results indicate that vemurafenib can promote the expression of EGF and activate the EGF/EGFR pathway. We also found removing vemurafenib from medium can significantly decreased *EGF* mRNA transcription (Fig. 3B). At the same time, we found that the mRNA and protein levels of *EGFR* were significantly increased in drug-resistant cells compared to the control (Fig. 3D, E).

**Figure 3.**
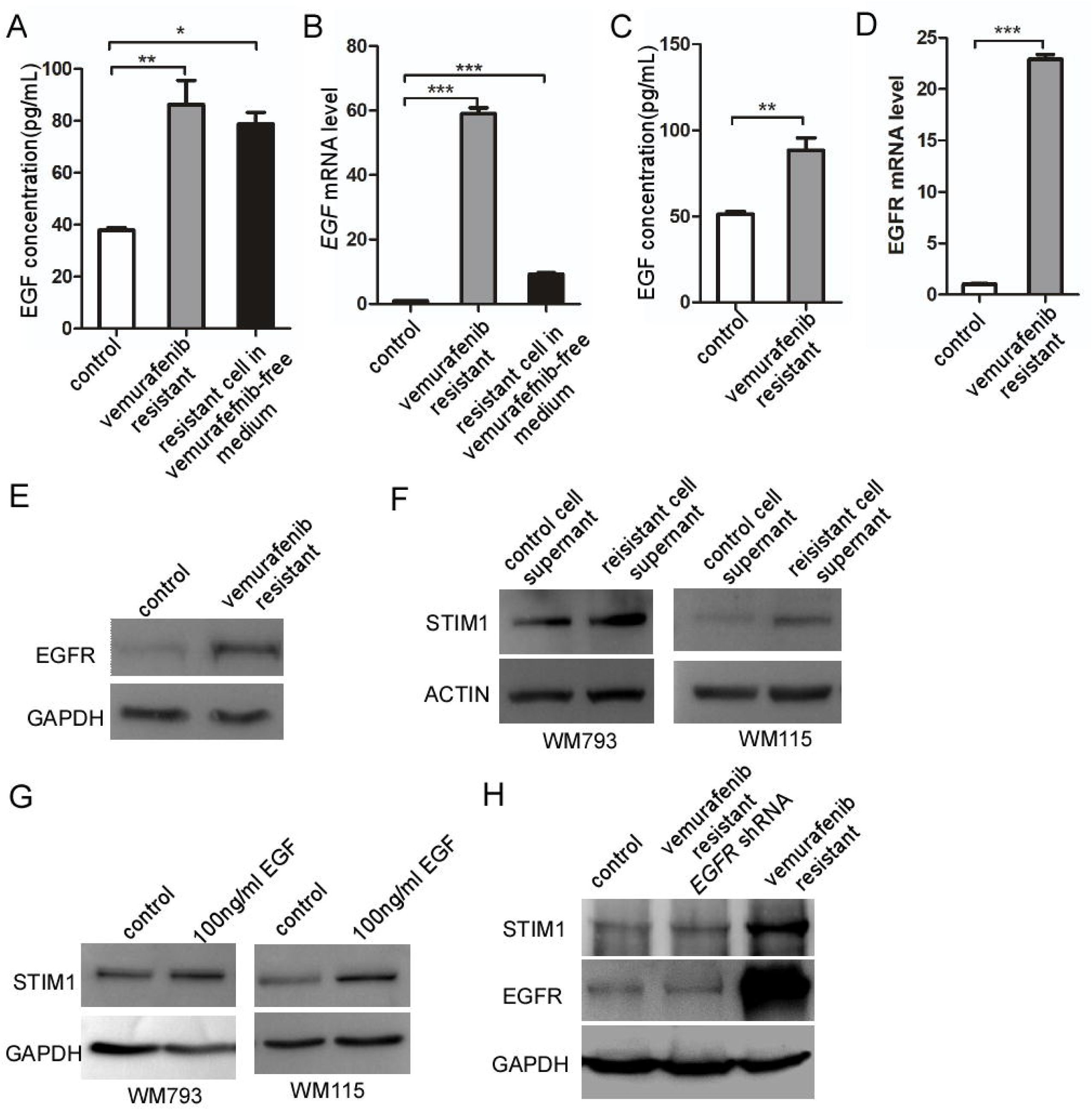
**Vemurafenib resistant induce STIM1 up-regulation through EGF/EGFR signaling** A. Elisa analysis of EGF concentration in control and vemurafenib resistant WM793 cells B. Q-PCR analysis of *EGF* level in control and vemurafenib resistant WM793 cells C. Elisa analysis of EGF concentration in control and vemurafenib resistant WM115 cells D. WB analysis of EGFR level in control and vemurafenib resistant WM793 cells E. Q-PCR analysis of *EGFR* level in control and vemurafenib resistant WM793 cells F. WB analysis of STIM1 level in WM793 and WM115 cells treated with control and vemurafenib resistant cell supernatant G. WB analysis of STIM1 level in WM793 and WM115 cells treated with control and EGF H. WB analysis of the effect of *EGFR* shRNA on vemurafenib resistance induced stim1 upregulation

To examine whether vemurafenib resistance promotes STIM1 expression through the EGF/EGFR pathway, we first collected supernatants from control and vemurafenib resistant cells and treated WM793 and WM115 cells. The results showed that the supernatant from vemurafenib resistant cells significantly increased STIM1 levels in both cell lines (Fig. 3F).

To further investigate whether vemurafenib resistance is regulated by EGF at STIM1 levels, we treated WM793 and WM115 cells with EGF, respectively. The results showed that EGF treatment significantly increased STIM1 levels in WM793 and WM115 cells (Fig. 3G), and *EGFR* shRNA significantly inhibited STIM1 levels in vemurafenib resistant cells (Fig. 3H). Our results indicate that vemurafenib resistance regulates STIM1 expression through the EGF/EGFR pathway.

### 4. EGF/EGFR regulates STIM1 expression through Hippo/YAP1

To investigate the mechanism that Vemurafenib resistance induce STIM1 up-regulation (To explore how EGF pathway regulates the STIM1 level), we first detected the level of downstream factors regulated by EGF/EGFR signaling.

It has been reported that BRAF inhibitor resistance could reactive the MAPK pathway, and recent studies have shown the reactivation of MAPK pathway in vemurafenib-resistant melanomas. However, p-ERK was significantly inhibited in our Vemurafenib resistant WM793 cells (Fig. 4A). The results suggest that the increase in STIM1 levels caused by vemurafenib resistance may not be through the MAPK pathway.

**Figure 4:**
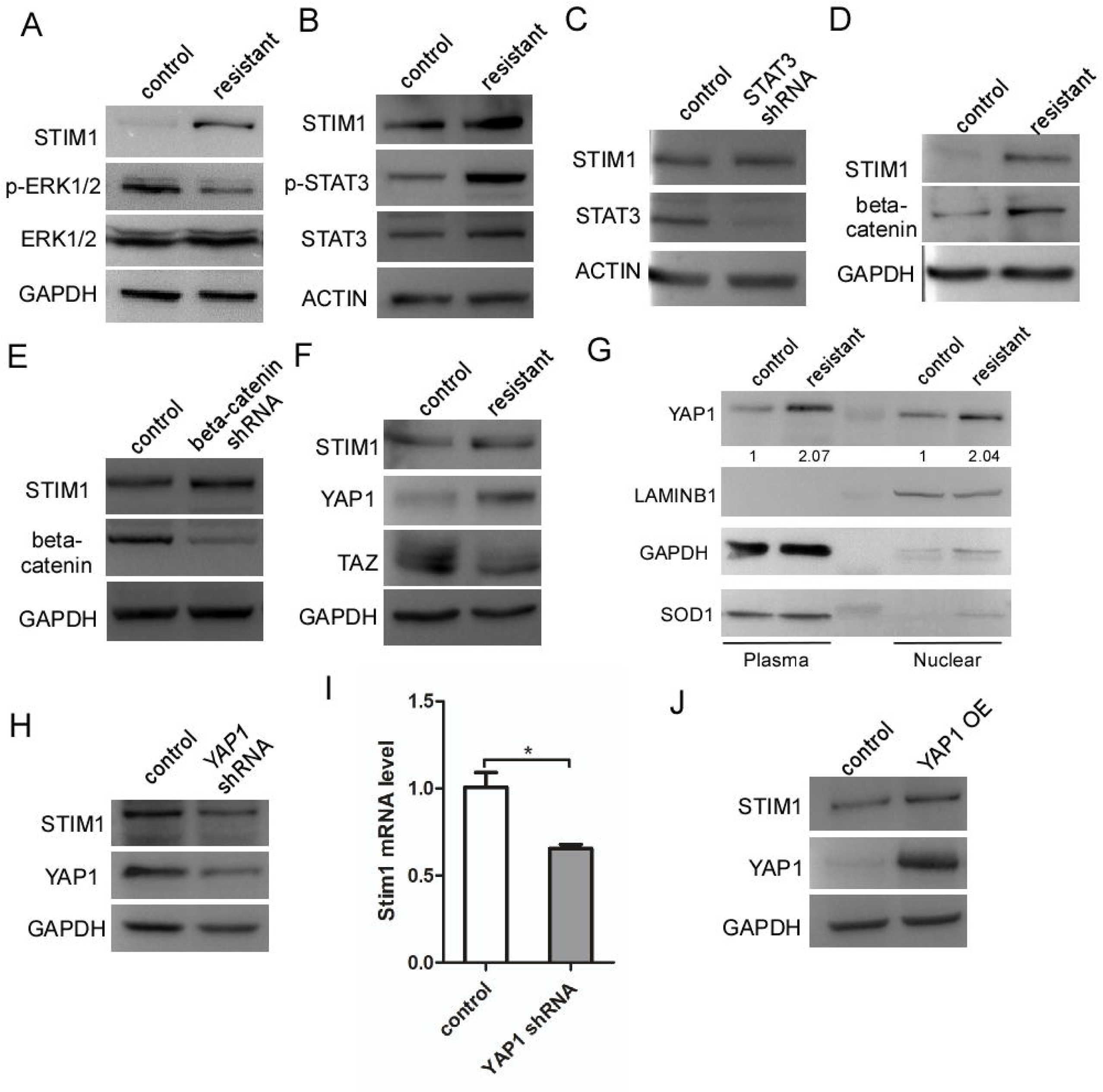
**BRAF inhibition increase STIM1 expression through YAP1** A. WB analysis of p-ERK level in control and vemurafenib resistant WM793 cells B. WB analysis of p-STAT3 level in control and vemurafenib resistant WM793 cells C. WB analysis of the effect of *STAT3* shRNA on STIM1 level D. WB analysis of β-catenin level in control and vemurafenib resistant WM793 cells E. WB analysis of the effect of β*-catenin* shRNA on STIM1 level F. WB analysis of YAP1 and TAZ level in control and vemurafenib resistant WM793 cells G. The effect of Braf resistance on YAP1 nucleus localization H. WB analysis of the effect of *YAP1* shRNA on STIM1 level I. Q-PCR analysis of the effect of *YAP1* shRNA on STIM1 level J. WB analysis of the effect of YAP1 overexpression on STIM1 level

*STAT3* is a transcription factor that plays central roles in various physiological processes and its deregulation results in serious diseases including cancer. STAT3 is upregulated in many cancers and activated via both EGFR-dependent and independent pathways[17], it is recently reported that IL6/STAT3 axis mediates resistance to BRAF inhibitors PLX4032(vemurafenib) in thyroid carcinoma cells[18]. Inhibition of STAT3 enhances vemurafenib sensitivity in colon cancers and melanoma harboring the *BRAFV600E* mutation[19, 20]. Interestingly, our RNA-seq results showed that IL6 was upregulated in vemurafenib resistant WM793 cells (Fig2. C), and the STAT3 level was significantly activated (Fig4. B), suggesting that IL6/STAT3 signaling was activated in vemurafenib resistant WM793 cells. However, our result showed that *STAT3* shRNA did not decrease STIM1 level in vemurafenib resistant WM793 cells (Fig4. C), which indicates that IL6/STAT3 does not mediate the regulation of STIM1 by BRAF inhibitors vemurafenib resistance.

Wnt/β-catenin signaling is frequently activated in tumorigenesis and contributes to the development of drug resistance. EGFR regulates β*-catenin* transactivation through a PKM2-dependent mechanism [21]. Our results show that β-catenin protein levels in resistant cells increased significantly, however β*-catenin* shRNA did not reduce the STIM1 level in vemurafenib resistance cells (Fig4. D-E), suggesting that WNT/β-catenin does not participate in the regulation of STIM1 by vemurafenib resistance.

It has been reported that EGF treatment of immortalized mammary cells triggers the rapid translocation of YAP into the nucleus along with YAP dephosphorylation[22]. It has also been shown that actin remodeling endows melanoma cells BRAF inhibitors resistance through YAP/TAZ activation[23]. To verify whether vemurafenib drug resistance is regulated by the EGF/EGFR **-**Hippo/YAP1 axis to regulate STIM1 levels, we first examined the level and nuclear uptake of YAP1. We found that the level of YAP1 was significantly increased in vemurafenib resistant cells, but not TAZ (Fig. 4F), and the nuclear localization of YAP1 in vemurafenib resistant cells was also significantly increased (Fig. 4G).

To verify the role of YAP1 in regulating STIM1 levels, we first knocked-down YAP1 in vemurafenib resistant cells. *YAP1* shRNA significantly decreased STIM1 protein and mRNA levels (Fig. 4H-I), and ectopic expression of YAP1 significantly increased STIM1 level (Fig. 4J). Our results suggest that Hippo/YAP1 mediates the regulation of STIM1 by the EGF/EGFR pathway.

In summary, vemurafenib resistance can activate multiple pathways related to tumor recurrence and metastasis, but only Hippo/YAP1 may be responsible for STIM1 upregulation

### 5. The YAP1/TEAD2 axis-mediated BRAF inhibitor resistance induced STIM1 up-regulation

*YAP1* is a co-activator of the transcription factor *TEAD* family, and it interacts with *TEAD* to regulate gene expression. To understand the molecular mechanisms by which *YAP1/TEAD* regulates *STIM1* transcription. We first analyzed the difference in the level of TEAD in WM793 control and vemurafenib resistant cells.

Interestingly, we found that the expression level of TEAD2 in vemurafenib resistant cells was significantly increased (Fig. 2C), indicating that TEAD2 might participate in the increase in the level of STIM1 caused by vemurafenib resistance. To investigate if vemurafenib induced STIM1 upregulation through YAP1/TEAD2, we first ectopically expressed TEAD2 in WM793 cells, and the result showed that TEAD2 overexpression can significantly increase STIM1 levels (Fig. 5A). We then confirmed the result through knocking down TEAD2 in WM793 control vemurafenib resistant cells respectively. We found that the STIM1 protein level dramatically decreased after TEAD2 knockdown (Fig. 5B-C).

**Figure 5.**
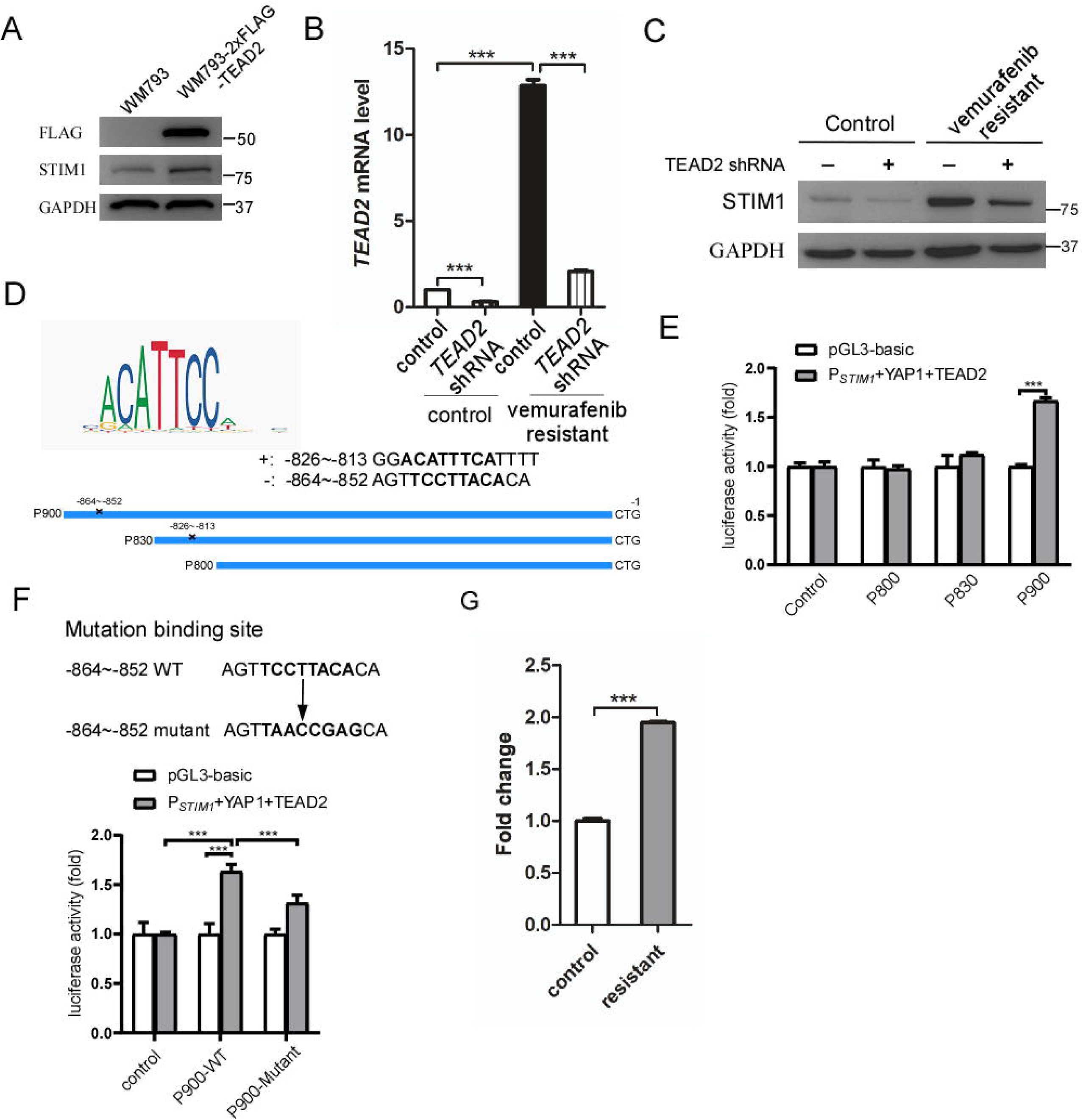
**YAP1/TEAD2 directly regulate Vemurafenib resistance induced STIM1 upregulation** A. WB analysis of the effect of *TEAD2* overexpression on STIM1 level. B. Q-PCR analysis of the effect of *TEAD2* shRNA efficiency. C. The effect of *TEAD2* shRNA on *STIM1* mRNA level. D. Motif of *TEAD2* transcription factor on the top left, and graphical representation of predicted binding sites. Forecasting tool supplied by JASPAR website (https://jaspar.genereg.net/). E. Luciferase activity of promoter fragments in contrast to pRL-TK in control and *YAP1-TEAD2* overexpression cell lines. F. Luciferase activity of the P-900 wild type and mutant in contrast to pRL-TK in control and *YAP1-TEAD2* overexpression cell lines. G. CHIP analysis of the binding of YAP1 to the STIM1 promoter in WM793 control and vemurafenib resistant cells. ***p<0.001.

To understand the molecular mechanisms by which YAP1/TEAD2 regulates vemurafenib resistance-mediated *STIM1* transcription, we examined the *STIM1* promoter for TEAD2 binding elements and identified two potential TEAD2 binding sites at -864∼-852 and -826∼-813 (Fig. 5D). To determine whether these two potential TEAD2 binding elements were required for TEAD2 to increase *STIM1* transcription, we use P900, P830 and P800 luciferase reporter and performed dual luciferase assay to investigate the TEAD2 binding site, the result showed that only P900 luciferase activity increase significantly when co-expression YAP1 and TEAD2, indicating that TEAD2 binds to -864∼-852 site in the *STIM1* promoter (Fig. 5D, E). To confirm the result, we mutated -864∼-852 core binding sites and repeated the dual luciferase assay, and found that the luciferase activity was significantly reduced after mutation of - 864∼-852 core binding sites (Fig 5F).

We then employed the CHIP assay using the YAP1 antibody, due to the limitations of the TEAD2 CHIP antibody. The result showed that the YAP1 antibody could successfully pulldown more *STIM1* promoters between -831 bp∼-900 bp in vemurafenib resistant cells than normal cells (Fig. 5G).

These results suggest that YAP1/TEAD2 mediates the transcription of *STIM1* by vemurafenib resistance. In addition, our present study revealed that vemurafenib-induced STIM1 upregulation through activation of the EGF/EGFR-YAP1/TEAD2 axis.

### 6. The YAP1/TEAD2-STIM1 axis is associated with melanoma progression

To evaluate the clinical significance of the YAP1/TEAD2-STIM1 axis in melanoma progression, we first analyzed the correlation between YAP1 and STIM1 levels in the TCGA database. The result indicated that YAP1 was positively associated with STIM1 levels (Fig. 6A). We then examined the levels of YAP1 and STIM1 in melanoma tissues and investigated the correlation between YAP1 and STIM1 in melanoma progression. All 180 cases in TMA slides were stained with anti-YAP1 and STIM1 antibodies. Immunohistochemical results of YAP1 and STIMI1 were assessable in 67 cases, which were categorized into low/none, medium, and high according to the intensity of staining (Fig. 6B), and the statistical results showed a positive correlation between the immunohistochemical expression of YAP1 and STIM1 (P < 0.000, R = 0.47, Fig. 6C).

**Figure 6.**
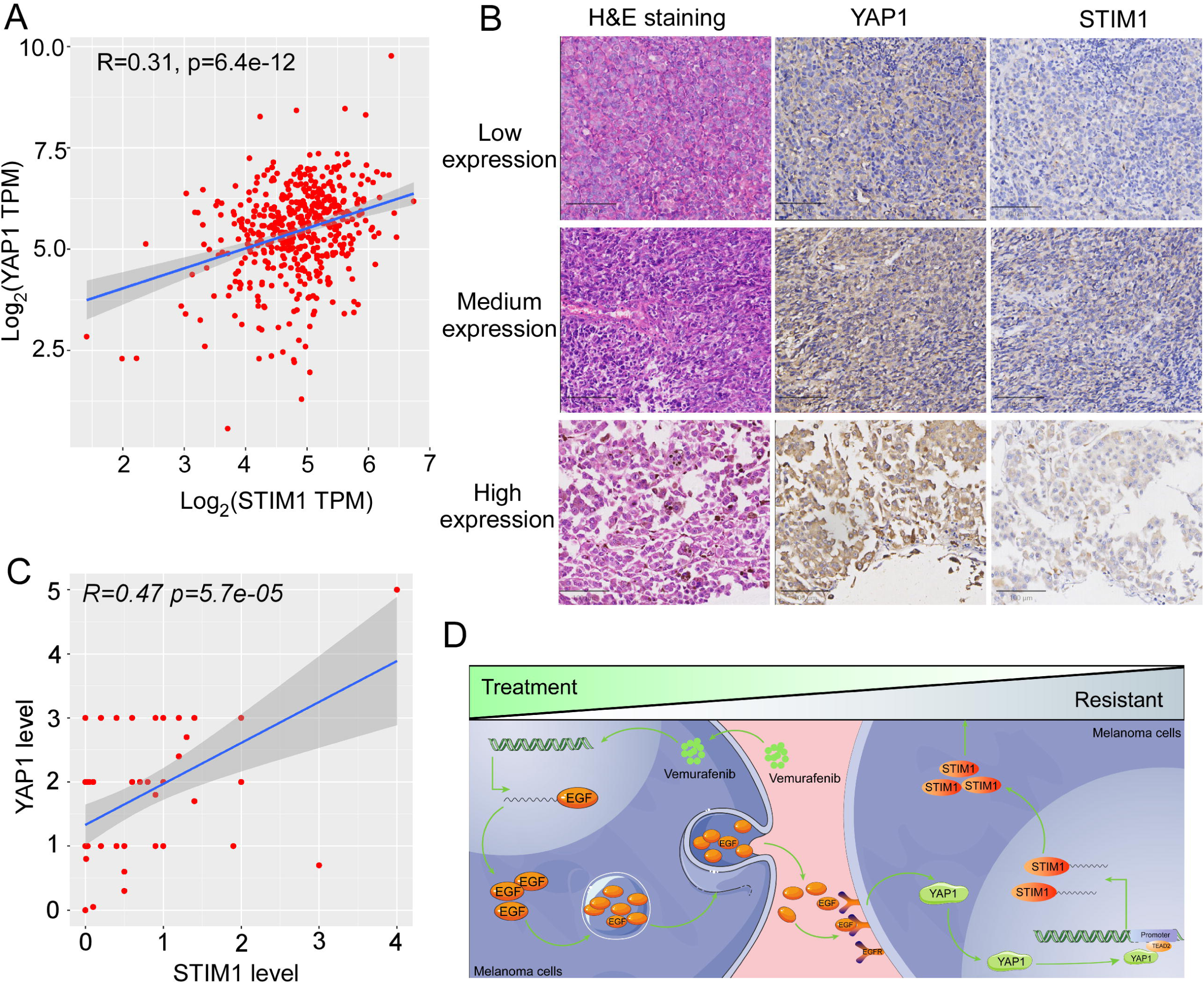
**Expression of YAP1 and STIM1 in melanoma tissues and survival analysis in melanoma patients.** A. Correlation analysis of *YAP1* and *STIM1* mRNA level from TCGA database. B. Representative images of YAP1 and STIM1 immunohistochemical staining in melanoma. C. The correlation analysis between YAP1 and STIM1 in melanoma /Correlation analysis of YAP1 and STIM1 level in patients with melanoma/ Comparison analysis of YAP1 and STIM1 expression levels in melanoma with different Clark grade. D. Model for vemurafenib resistance induced STIM1 upregulation.

In conclusion, our result indicated that vemurafenib treatment/drug resistance will lead to the increase of EGF and EGFR levels in melanoma. The activated EGF/EGFR pathway will promote YAP1 to enter the nucleus and then activate Hippo/YAP1 pathway. *YAP1/TEAD2* will bind to the *STIM1* promoter region and promote the transcription and expression of *STIM1*(Fig. 6D).

## Discussion

The *BRAF* mutation is the most prevalent genetic alteration in human melanoma, with more than 50% of the patients harboring the *BRAF(V600E)* mutation[3]. The discovery of mutant specific BRAF inhibitors, such as vemurafenib, has remarkably improved the survival of melanoma patients with oncogenic BRAF mutations[24]. However, vemurafenib resistance is still the main obstacles for the treatment of melanoma, and the mechanism of vemurafenib resistance induced melanoma invasion and metastasis is still not completely understood[25].

STIM1, the key sensor for SOCE channels, has been reported in tumorigenesis and metastasis[11, 12]. However, there are relatively few reports on the role of STIM1/SOCE in the occurrence and metastasis of melanoma. We have reported that SOCE-induced invadopodia formation leads to melanoma metastasis[11, 15]. Moreover, we found that STIM1 level is upregulated in BRAF inhibitor vemurafenib resistant cells[14], suggesting that STIM1 may respond to vemurafenib treatment. However, the role and regulatory mechanism of STIM1 in vemurafenib resistance is unknown. Ziyi Yang reported that the STIM1 expression level was related to the acquired resistance to imatinib in gastrointestinal stromal tumors[13]. Our recent research showed that BRAF inhibitor vemurafenib resistance induces STIM1 up-regulation, indicating STIM1 may respond to the vemurafenib resistance[14]. However, the mechanism of STIM1 upregulation by vemurafenib resistance is unclear. In the present study, we identified that the EGF/EGFR-YAP1/TEAD2 axis mediated STIM1 up-regulation in vemurafenib resistant melanoma cells. Vemurafenib resistance can lead to the increase of EGF and EGFR levels and activate the EGFR signaling pathway, which will promote the YAP1 nuclear localization, and then increase the expression of STIM1.

It has been previously reported that YAP1/TAZ activation confers BRAF inhibitor resistance in melanoma cells [23]. We also showed that STIM1 knockout significantly inhibited/reduced cell proliferation in the presence of vemurafenib. Future investigations on the mechanism by which STIM1 confers vemurafenib resistance is warranted.

In summary, our present study found that the EGF/EGFR-YAP1/TEAD2-STIM1 axis respond to vemurafenib therapy and resistance. Our finding not only demonstrated the mechanism by which vemurafenib resistance promotes STIM1 expression, but also suggested that targeting the EGF/EGFR-YAP1/TEAD2-STIM1 axis could improve the therapeutic efficiency of BRAF inhibitors in melanoma patients.

## Materials and methods

### Cell culture and inhibitor treatment

Melanoma cell line WM793 and WM115 were cultured in RPMI1640 medium supplemented with 10% fetal bovine serum (FBS) and 1% penicillin/streptomycin. Vemurafenib resistant WM793 and WM115 cells were made by adding 2 μM vemurafenib in RPMI1640 medium for 2 weeks and were maintained in RPMI1640 medium supplemented with 5% FBS and 1 uM vemurafenib. For the inhibitor treatment, cells (5 × 10^5^) were seeded in 6 well plates and treated with 2 μM vemurafenib or PF562711 for 12 h, 24 h, and 36 h after the cells attached to the dish.

### Western blot analysis

Western blotting was performed as described previously[26, 27]. Briefly, 50 μg protein of whole cell lysate from each sample was loaded on a 10% PAGE gel; the membrane was blocked in 5% non-fat milk in 1 × Tris-buffered saline (pH 7.4) containing 0.05% Tween-20 and subsequently probed with primary antibodies at a concentration of 1:1000 or 1: 5000 (GAPDH). The secondary antibodies were used at a concentration of 1:10,000 to 1:20,000. The proteins were visualized with the ECL-plus western blotting detection system (Tanon-5200Multi).

The following antibodies were used in this study: mouse monoclonal GAPDH(G8795; Sigma-Aldrich, Shanghai, China), mouse monoclonal STIM1(sc-166840; Santa Cruz, Shanghai, China), rabbit polyclonal Actin(sc-1616; Santa Cruz, Shanghai, China), mouse monoclonal Raf-B(sc-5284; Santa Cruz, Shanghai, China), mouse monoclonal EGFR(sc-373746; Santa Cruz, Shanghai, China), mouse monoclonal GAPDH(sc-32233; Santa Cruz, Shanghai, China), mouse monoclonal p-ERK(sc-7383; Santa Cruz, Shanghai, China), mouse monoclonal ERK(sc-514302; Santa Cruz, Shanghai, China), mouse monoclonal p-stat3(sc-8059; Santa Cruz, Shanghai, China), rabbit polyclonal β-catenin(A01211-40; Genscript, China), rabbit polyclonal YAP1(A1002; Abclonal), mouse monoclonal TAZ(sc-518026; Santa Cruz, Shanghai, China), rabbit polyclonal α-H3(ab1791; Abcam), mouse monoclonal Flag(F1804; Sigma-Aldrich), rabbit polyclonal STAT3(06596; Sigma-Aldrich)

### Luciferase Assay

The Dual Luciferase Assay was performed according to a previously reported protocol with minor modification[28]. The STIM1 promoter luciferase reporter constructs were generated by inserting 900bp, 830bp or 800bp human STIM1 promoter into pGL3 basic vector (Promega) between XhoI and HindIII. To perform dual luciferase reporter assay, 5,0000 293T cells were seeded in 12-well plates and cultured overnight. Cells were transfected with 1 μg/well STIM1 promoter reporter together with 100 ng/well Renilla luciferase construct (Prl-TK) using Lipofectamine 2000. 24h after transfection, the cells were was lysed and Cell lysates were subjected to dual reporter luciferase assays according to the manufacturer’s instructions (Promega).

### Tissue microarray and immunohistochemistry

Tissue microarrays (TMAs) were constructed using a manual tissue microarray instrument (Beecher Instruments) equipped with a 2.0 mm punch needle, as we previously described[14]. The levels of STIM1 and YAP1 were assessed via the average of 5 count fields per patient in the original magnification of X200 on light microscopy. The expression of STIM1 and YAP1 was scored based on staining intensity. Staining intensity was subclassified as follows: 0, negative; 1, weak; 2, moderate; and 3, strong.

The primary antibodies used in this study were as follows:

STIM1, Rabbit #3113 CST IHC: 1:200

YAP1, Rabbit #PA5-40327 Thermo Fisher Tech. IHC: 5 µg/mL

## Data availability

All data are available in the manuscript or the supplementary materials.

## Statistical analysis

All the experiments were repeated three time and each experiment was performed in 3 replicates per sample. Data were analyzed using SPSS 19.0 and GraphPad Prism 6.0. Student’s t-test, Spearman correlation, Kaplan-Meier, log-rank test and Cox regression survival analysis were according to a previous report[29] and Statistical significance was defined as \**P*[<[0.05, \*\**P*[<[0.01 or \*\*\**P*[<[0.001.

## Acknowledgements

We thank Dr. Shengyu Yang and Jing Li for critical reading of the manuscript. We especially thank Prof. Baowei Jiao for *YAP1* and *EGFR* shRNA plasmid.

